# Two-Photon Lithography-Fabricated Deterministic Lateral Displacement Microfluidic System for Efficient Minicell Purification in Cancer Therapy

**DOI:** 10.1101/2025.06.19.660580

**Authors:** Sharaj Hegde Sharavu, Sagar Bhagwat, Büsra Merve Kirpat Konak, Barbara Di Ventura, Pegah Pezeshkpour, Bastian E. Rapp

## Abstract

Chromosome-less minicells, derived from aberrant polar division events of bacterial cells, have emerged as promising nanocarriers for targeted cancer drug delivery due to their unique characteristics. A major challenge in their purification process lies in effectively isolating such spherical minicells (<1 µm) from their rod-shaped parental cells (1-10 µm). This study investigates the use of Deterministic Lateral Displacement (DLD) microfluidic systems for minicell purification, leveraging Two-Photon Lithography (TPL) for the rapid prototyping of high-resolution designs optimized for this purpose. Under laminar flow conditions, we investigated key DLD design parameters including symmetric and asymmetric post gaps, outlet widths, dual post arrays, fluidic-resistance-optimized design. To enhance separation efficiency, we developed a two-stage microfluidic separation system combining a spiral inertial chip and an optimized DLD chip in series. Utilizing high-resolution TPL for chip fabrication of an inertial chip with 12 spirals and an asymmetric DLD chip with a 2 µm downstream post gap, we achieved a separation efficiency of 94%. This high efficiency achieved using microfluidics for the separation of cells differing in both shape and size, demonstrates the potential of advanced microfluidic systems in cell sorting.

**TOC:** This study presents the use of Deterministic Lateral Displacement (DLD) microfluidic systems, fabricated via high-resolution 3D printing, Two-Photon Lithography, to isolate minicells for targeted cancer drug delivery. By optimizing post geometries, array designs, fluidic resistance, and integrating a spiral inertial stage, the system achieves 94% separation efficiency. These advancements ensure high throughput, operational stability, and continuous sorting of bacterial cells.

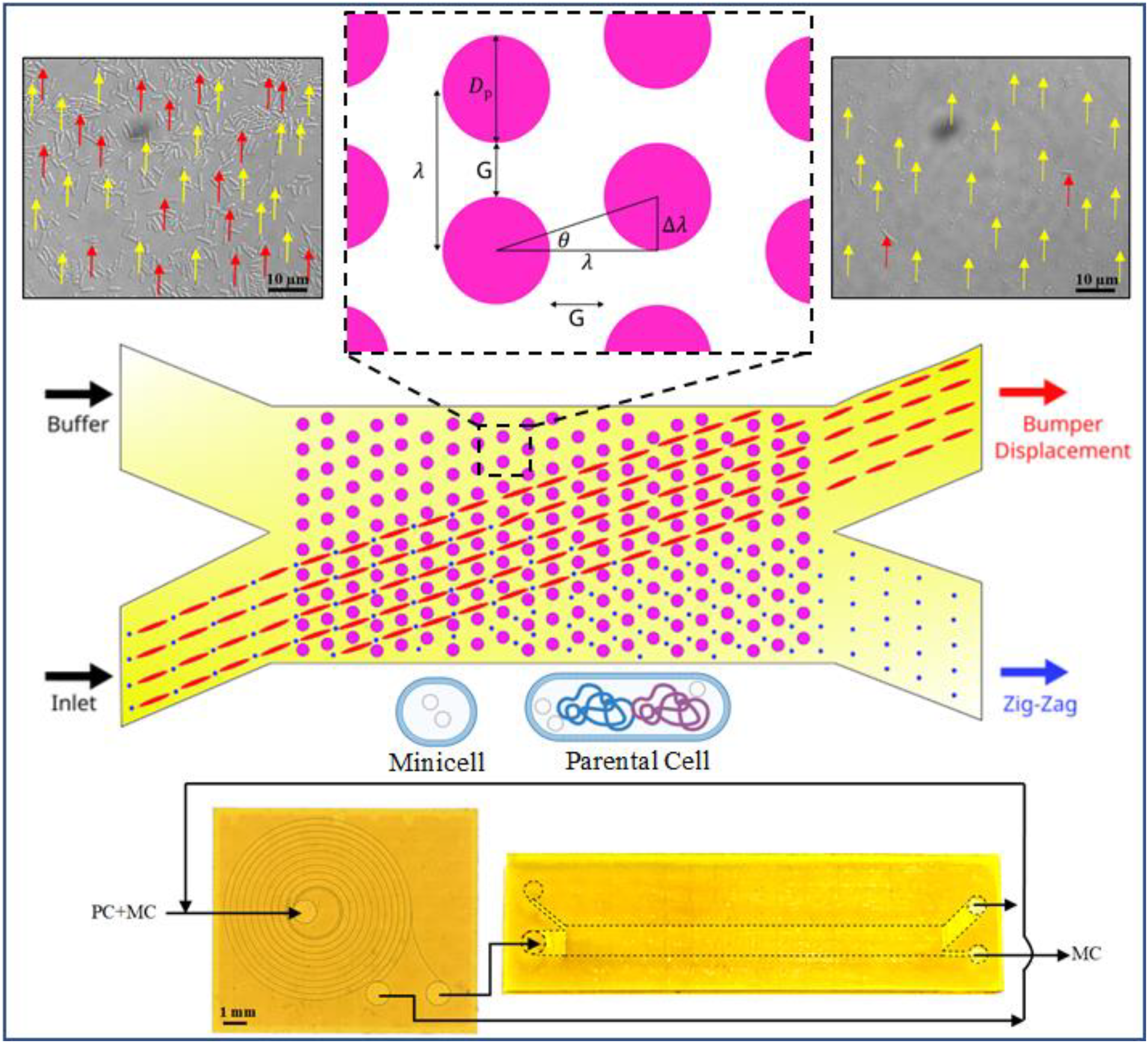

## 1. Introduction

Cancer remains one of the most lethal diseases of the 21^st^ century, claiming millions of lives annually.^[1]^ Chemotherapy is among the most widely used cancer treatments; however, despite its effectiveness, it suffers from severe side effects caused by damage to healthy cells, due to its systemic administration. The concept of targeted cancer therapy –where the drugs are delivered directly to the cancer cells, sparing healthy ones– emerged in the late 80s and has achieved some clinical success.^[2–4]^ Various approaches fall within this category, one of which involves the use of nanoparticles (NPs) to deliver the chemotherapeutic drugs. These nanoscale carriers are larger than individual drug molecules but remain within the nanometer range. NPs, into which chemotherapeutic drugs are encapsulated, accumulate in the cancer microenvironment either passively due to the enhanced permeability and retention (EPR) effect or actively via transport and retention (ATR).^[5,6]^ NPs are typically equipped with targeting moieties—such as chemical compounds, peptides, or proteins—that specifically recognize and bind to surface receptors overexpressed on cancer cells, facilitating selective drug delivery to particular cell types. An emerging type of NPs is bacterially derived minicells.^[7]^ Minicells arise from asymmetric cell division events, resulting in a small round cell devoid of chromosomal DNA, and a parental cell^[8]^. Compared to other inorganic or organic NPs, minicells have several advantages: their production does not require harsh chemicals, they are very robust (they do not rupture, nor do they leak their contents), and they can be easily genetically manipulated to deliver biologics, such peptide/proteins or nucleic acids. While attenuated bacteria share these features with minicells and are indeed applied in cancer therapy^[9]^, the lack of the chromosome greatly reduces safety concerns for therapeutic applications in humans. Modifying their surfaces with targeting antibodies allows minicells to specifically recognize and interact with cancer cells, thereby enhancing treatment efficacy while reducing off-target effects.^[10–12]^

For therapeutic applications, it is paramount to eliminate parental cells from minicells reaching a purity of 1-2 parental cells in 10^8^-10^9^ minicells. Minicells are typically separated from parental cells using a series of centrifugation and filtration steps.^[13,14]^ Moreover, the purification protocol most often includes an antibiotic treatment targeting specifically the growing parental cells.^[14]^ This purification procedure is time-consuming, tedious and costly. We set ourselves the goal to develop a novel minicell purification method based on microfluidics.^[15–17]^

Microfluidic methods for particle separation are generally categorized into active^[18–21]^, involving real-time manipulation of particle movement, and passive techniques^[22]^, the latter including hydrodynamic filtration^[23]^, pinched flow fractionation^[24]^, viscoelastic separation^[25]^, inertial microfluidics^[26]^ and deterministic lateral displacement (DLD) separation.^[27]^ We selected DLD microfluidics due to its potential for high separation efficiencies.^[28]^ This method enables precise and deterministic sorting of particles based on intrinsic properties like size^[29]^, shape^[30]^, and deformability. Separation is achieved using engineered microfluidic structures to guide particles along deterministic trajectories through arrays of pillars.^[31,32]^ In a DLD array, particles larger than a critical size follow a distinct lateral displacement path, while smaller particles travel along the fluid streamlines in a zigzag pattern.^[33]^ In our case, fabrication of DLD microfluidic chips is particularly challenging due to the extremely small microscale features required to separate submicron-sized minicells.

Our previous designs based on DLD achieved a good degree of separation.^[30,34]^ Yet, this purity is not sufficient for downstream therapeutic applications. A key challenge in a procedure based on microfluidics where the starting material is a mixed population of minicells and parental cells which did not undergo any treatment (such as a first centrifugation step to roughly separate parental cells from minicells and an antibiotic treatment to elongate and lyse parental cells) is the presence of very small parental cells.

In this work, we develop a new microfluidic platform leveraging two-photon lithography (TPL) for rapid prototyping of different DLD designs. Indeed, microfabrication techniques such as photolithography^[35]^, stereolithography^[36]^, soft lithography^[37]^ and TPL^[37]^ facilitate the creation of a wide range of microsystems. Among all, TPL-based fabrication technique allows for high-resolution rapid prototyping of multiple design iterations.^[34]^ Using focused femtosecond laser pulses, TPL induces localized polymerization within photosensitive materials, enabling the fabrication of intricate 3D structures with resolutions down to the nanoscale.^[38–41]^

We demonstrate a combined spiral inertial and DLD microfluidic chip design achieving separation efficiencies of up to 94 % in minicell separation at throughputs of 5 µL/min. Despite affording an efficiency below the desired one of >99%, this design represents a stark improvement compared with previous designs.

These novel microfluidic systems provide a versatile platform for cell separation and manipulation, enabling precise control over biological samples for a variety of applications beyond minicell purification.

## 2. Results and Discussions

### 2.1 Minicell production

To trigger minicell formation, we used a previously described method^[42]^ that relies on the overexpression of the MinDΔ10 protein, which disrupts the normal function of the Min system responsible for positioning the division plane to mid-cell. Briefly, *Escherichia coli* BL21 cells were transformed with a plasmid encoding the blue light-responsive transcription factor BLADE, which drives the expression of MinDΔ10. MinDΔ10 is unable to bind to the membrane, but interacts with MinC, leading to its accumulation in the cytoplasm where it cannot counteract FtsZ. Consequently, FtsZ is free to form at the poles, giving rise to minicells (**Figure 1a**). Bacterial cells additionally expressed a fluorescent protein for visualization under a fluorescence microscope. Microscopic analysis of the cell population after blue light illumination showed a mixture of minicells and parental cells, some of which were as short as 1-2 μm (**Figure S1**).

**Figure 1.**
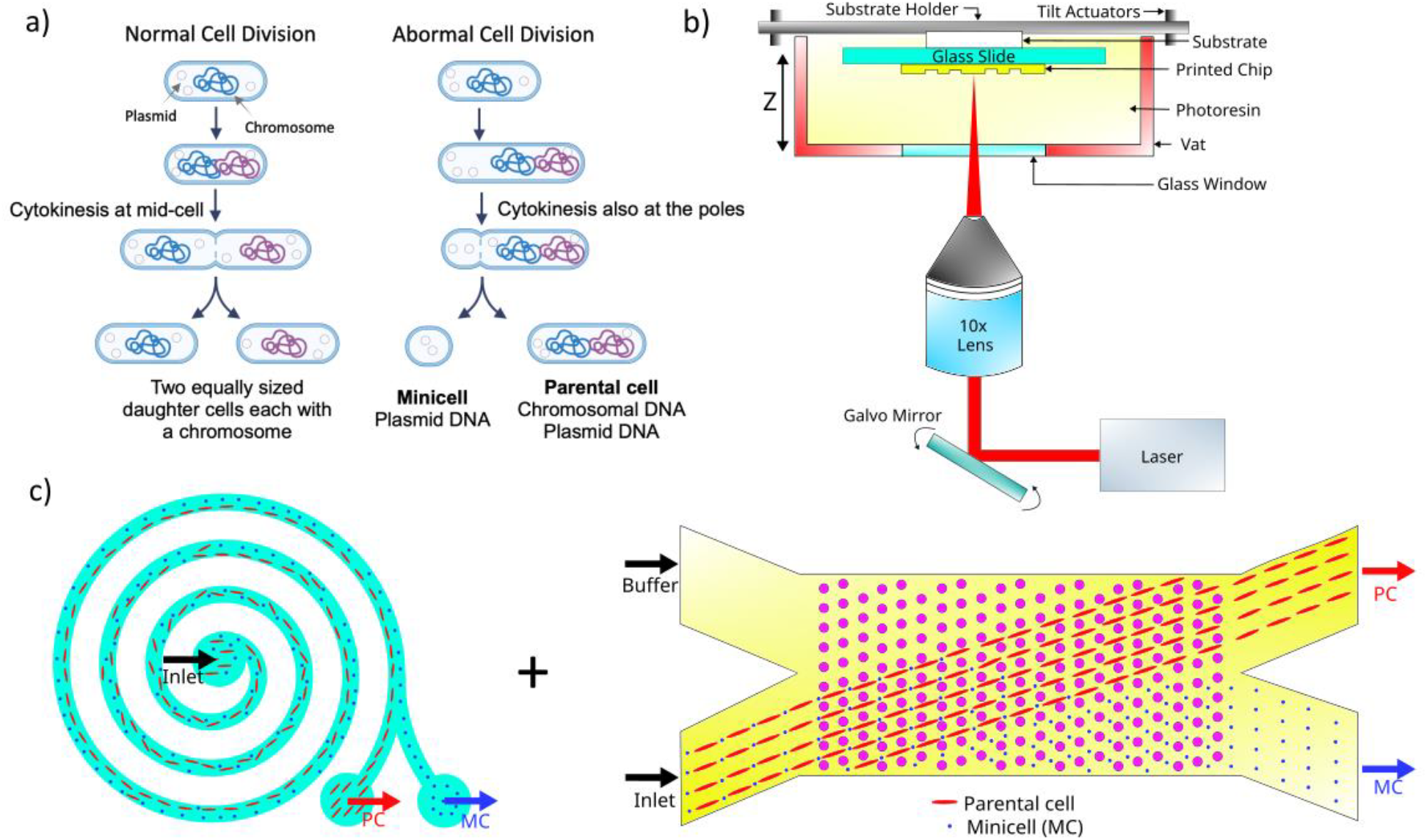
Schematics for minicell purification. a) Scheme of normal and abnormal bacterial cell division leading to minicell formation (Created with BioRender) b) TPL fabrication in vat mode for the fabrication of microfluidic chips c) Schematics of the spiral inertial chip as well as the DLD chip used in combination. PC, parental cell. MC, minicell.

### 2.2 TPL & DLD parameters

We started by optimizing the key parameters of the DLD design that influence microscale flow mechanics within the chip, aiming to improve cell separation based on size and shape. We employed TPL to fabricate the microfluidic chips with submicron resolution (**Figure 1b**). A combination of microfluidic systems and design optimizations were investigated towards the goal of minicell purification (**Figure 1c**). Our previous studies showed that DLD post arrays can be used for shape based separation by passively displacing longer parental cells in a bumped path towards the top outlet, while the minicells follow a zigzag path through the downstream gaps, ultimately being extracted from the bottom outlet.^[30,34]^ DLD is one of the most effective passive microfluidic system for achieving high separation efficiency.^[28,43–50]^ The threshold size between these two displacement modes is determined using the critical diameter (D_C_), which is calculated using the DLD design parameters.^[51–53]^ The individual posts in the DLD array are repeated at distances of λ, which is the sum of gap (G) and the post diameter (D_P_) (Equation 1). Furthermore, each adjacent column is offset from the previous one by an angle (θ), corresponding to a distance (Δλ), and is represented by the post shift fraction (ε) (Equation 2).^[54]^

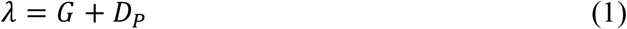

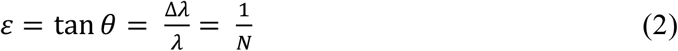

where N is the number of streamlines.

These parameters are used to calculate the critical diameter (D_C_) for which a particle enters zigzag displacement (Equation 3).^[54]^

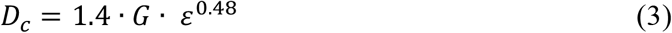

The parameters in the post array of the DLD design are shown in **Figure 2a**.^[28]^

**Figure 2.**
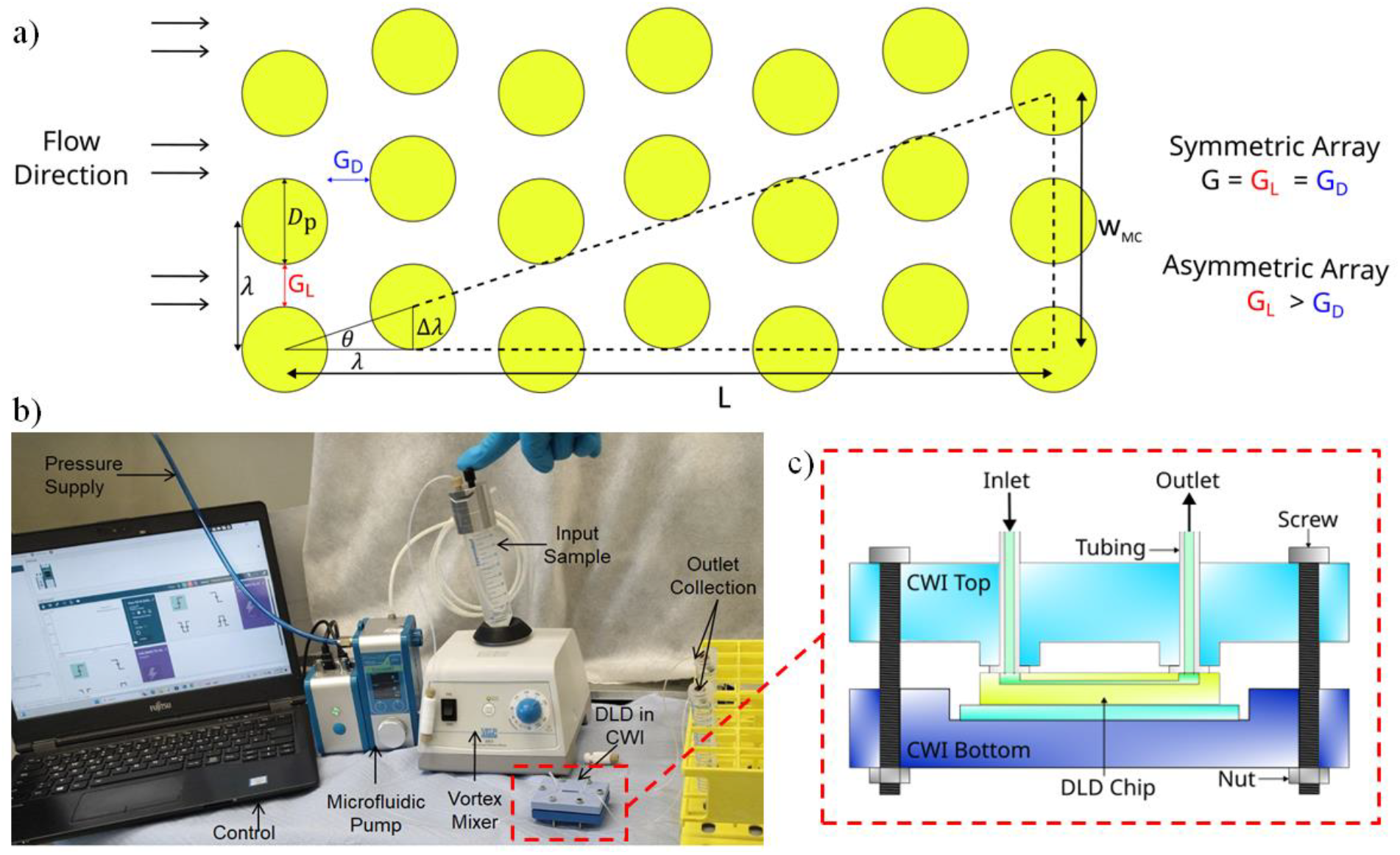
a) DLD parameters for the post array b) Image of the experimental setup c) Schematic of the DLD chip in CWI

The width of the minicell outlet is proportional to the total chip length and the shift angle. All DLD chips in this study have a channel length (L) of 19 mm. The width of the minicell outlet in the zigzag displacement collection region (*W*_*MC*_) can be determined from Equation 4:

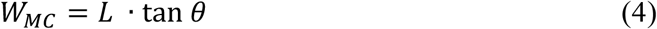

The devices were manufactured using TLP on a standard microscope slide, which was first plasma-cleaned and then surface-functionalized with 3-methacryloxypropyldimethylchlorosilane (MACS). The experimental setup for the minicell purification is shown in **Figure 2b** including the “Chip to World Interface” (CWI) as seen in **Figure 2c**.

The minicell separation efficiency (η) reflects the extent of minicell purification achieved with the microfluidic separation and is calculated comparing the ratio of parental cells to minicells before and after separation from the minicell outlet (Equation 5).^[30]^

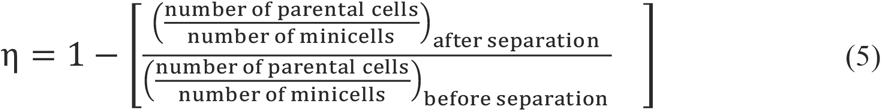

To optimize the design for the highest efficiency, we investigated different chip configurations including symmetric and asymmetric post gaps, minicell outlet width, fluidic optimized design, dual array, spiral inertial chip and a multi-stage separation system using both spiral inertial chip and DLD in series.

We used the TPL printer to print the DLD chip on a 38 mm × 25 mm functionalized glass substrate utilizing the commercially available UpPhoto photoresin. The posts were printed in high resolution ‘voxel’ mode, with each voxel measuring 730 nm in the XY plane and 9.2 µm in height along the Z axis, at a volumetric printing speed of 20 mm^3^/h. The remainder of the chip was printed using ‘simple’ mode to accelerate the process. The total printing time for the DLD chip and spiral inertial chips were approximately 12 and 8 hours, respectively. Post with a diameter of 25 µm and gaps between consecutive posts ranging from 5 µm to 2 µm were printed using TPL to evaluate the printing resolution of the complex array design (**Figure 3a)**. We found that gaps smaller than 2 µm tend to fuse while using the 10x objective, which is why larger gaps were chosen.

**Figure 3.**
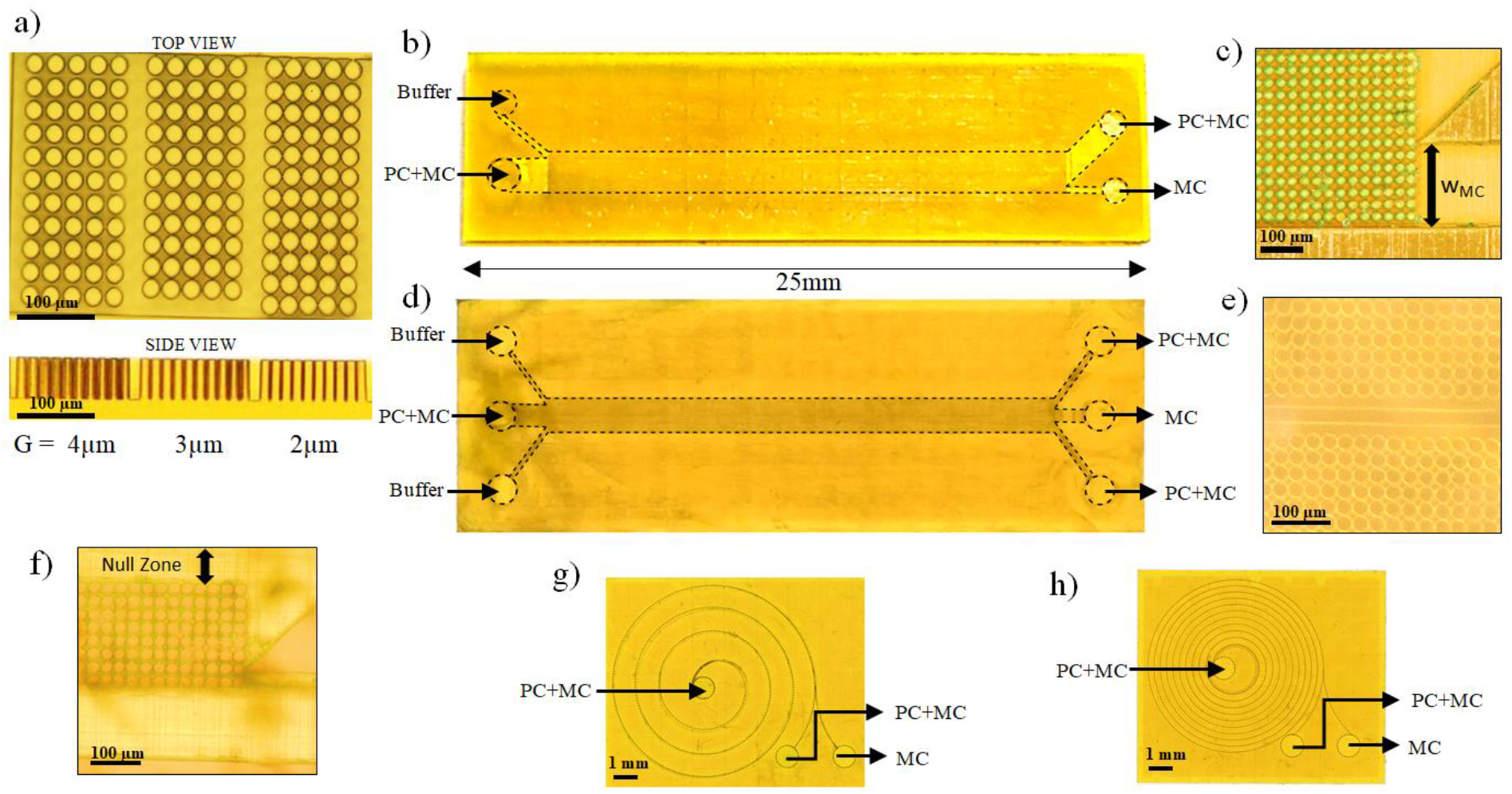
Optical microscope images a) TPL resolution of the top and side view for different gaps (scale 100 µm) b) Single array DLD. c) Minicell outlet width d) Dual array DLD. e) Mirrored posts in dual array DLD. f) Null zone in fluidic resistance optimized DLD g) 4-Spiral inertial chip. h) 12-Spiral inertial chip. b-h) PC, parental cell. MC, minicell.

### 2.2 DLD parameter variations and optimization

The standard single array DLD chip (**Figure 3b**) has a channel length of 19 mm, post diameter of 25 µm and post shift angle of 0.75°. Decreasing post gaps increased separation efficiency by lowering the critical diameter. As the post gap decreased from 7 µm to 4 µm, separation efficiency improved significantly from 36 % to 71 %, enabling more precise cell sorting (**Table S1**). Such increase in efficiency came at the cost of a reduced flow rate, which dropped from 29 µL/min to 8 µL/min.

The theoretical minicell outlet width (*W*_*MC*_), as shown in **Figure 3c** and calculated using Equation 4, is 0.25 mm. During the DLD process, there is a chance that parental cells may accidentally slip through the downstream gap and be displaced in the zigzag mode. This results in a transition zone, further reducing the optimal *W*_*MC*_. The chip was tested with different minicell outlet widths. A minicell outlet width of 0.3 mm, which is larger than the theoretical limit, resulted in an efficiency of only 48 %. We observed a significant increase in efficiency from 71 % to 80 % by reducing the minicell outlet width from 0.25 mm to 0.20 mm. By reducing this width, the likelihood of parental cells accidentally slipping into lower streamlines and being collected in the minicell outlet was substantially reduced. To increase the flow rate of the chip, we tested a dual array system that is mirrored with respect to the centerline (**Figure 3d**,**e**). This configuration effectively doubled the volume throughput of the chip while maintaining a separation efficiency of 75 %.

Next, we investigated asymmetric post arrays with lateral gaps (G_L_) larger than the downstream post gaps (G_D_). The incorporation of asymmetric post gaps can enhance efficiency while reducing the risk of clogging. We tested DLD chips with lateral post gaps increasing from 4 μm to 7 μm, while maintaining a fixed downstream gap of 4 μm. Both separation efficiency and flow rate improved as the lateral post gap increased, reaching a peak efficiency of 84 % at G_L_ = 6 μm. However, further increasing the lateral post gap beyond 7 μm reduced efficiency of 81 %, as more streamlines caused fewer minicells to follow zigzag displacement. Taken together, these observations indicate that increasing the lateral post gap in an asymmetric array enhances efficiency by preventing accidental zigzag displacement of parental cells. However, beyond a certain point, further increases inhibit minicell zigzag displacement, thereby reducing efficiency.

We additionally optimized the DLD design based on the fluidic resistance (**Figure 3f**). Reducing the fabrication complexity, this chip features only ten rows of posts in the lateral displacement region and a narrow “null zone” with no posts, added to lower the total fluidic resistance. The reduction of fluidic resistance enables the development of chips with smaller downstream post gaps which reached 91 % efficiency for post gaps of G_D_: G_L_ taken as 2 μm: 4 μm.

### 2.3 Spiral Inertial Chip and Multi-stage microfluidic platform

The spiral inertial chip design (100 µm channel height and width) is a common device in microfluidics allowing particle sorting based on shear gradient-induced lift force, wall-induced lift force and Dean forces which move the particles to equilibrium positions within the channel.^[55–58]^ We therefore investigated spiral chips with 4 spirals (**Figure 3g**) and 12 spirals (**Figure 3h**) for minicell purification. The Dean forces that are critical for separation depend on the flow rate: a low flow rate leads to ineffective separation, while a high flow rate causes excessive vorticity.

Experiments showed that an input pressure of 1000 mbar achieved a separation efficiency of 58 % for the 12-spiral chip and 32 % for the 4-spiral chip. Based on these results, we decided to combine the spiral inertial microfluidics chip and the DLD chip in a series connection to create a multi-stage microfluidic separation system (**Figure 4a**). In the first stage, the spiral inertial microfluidic chip utilizes inertial lift forces and Dean flow effects to remove cells of larger aspect ratio, reducing clogging risks in the downstream DLD chip thereby enhancing the deterministic displacement of particles. The experiments were performed with 1000 mbar inlet pressure for the spiral inertial chip and then passed through DLD chips with 2 µm : 3 µm and 2 µm : 4 µm post gaps, respectively. This multistage microfluidic system reached efficiencies as high as 94 % with minimal clogging of the DLD channels. **Figure 4b-h** summarizes all the design configurations, the correlated purification efficiency and the zoomed images of minicells and parental cells at the minicell outlet. The purification efficiency values achieved in this work are shown in **Table 1**.

**Table 1.**
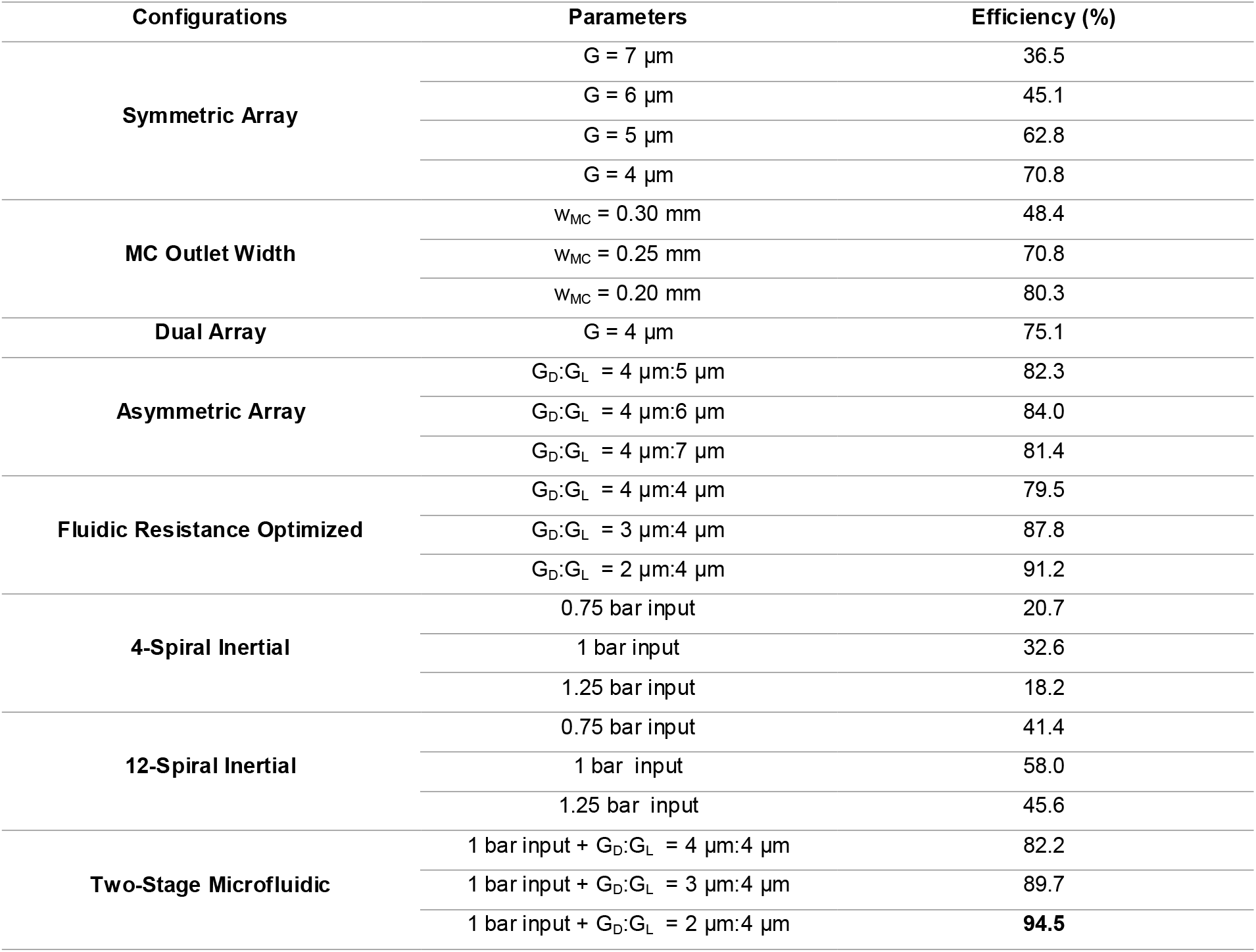
Efficiencies of all configurations tested in this study.

**Figure 4.**
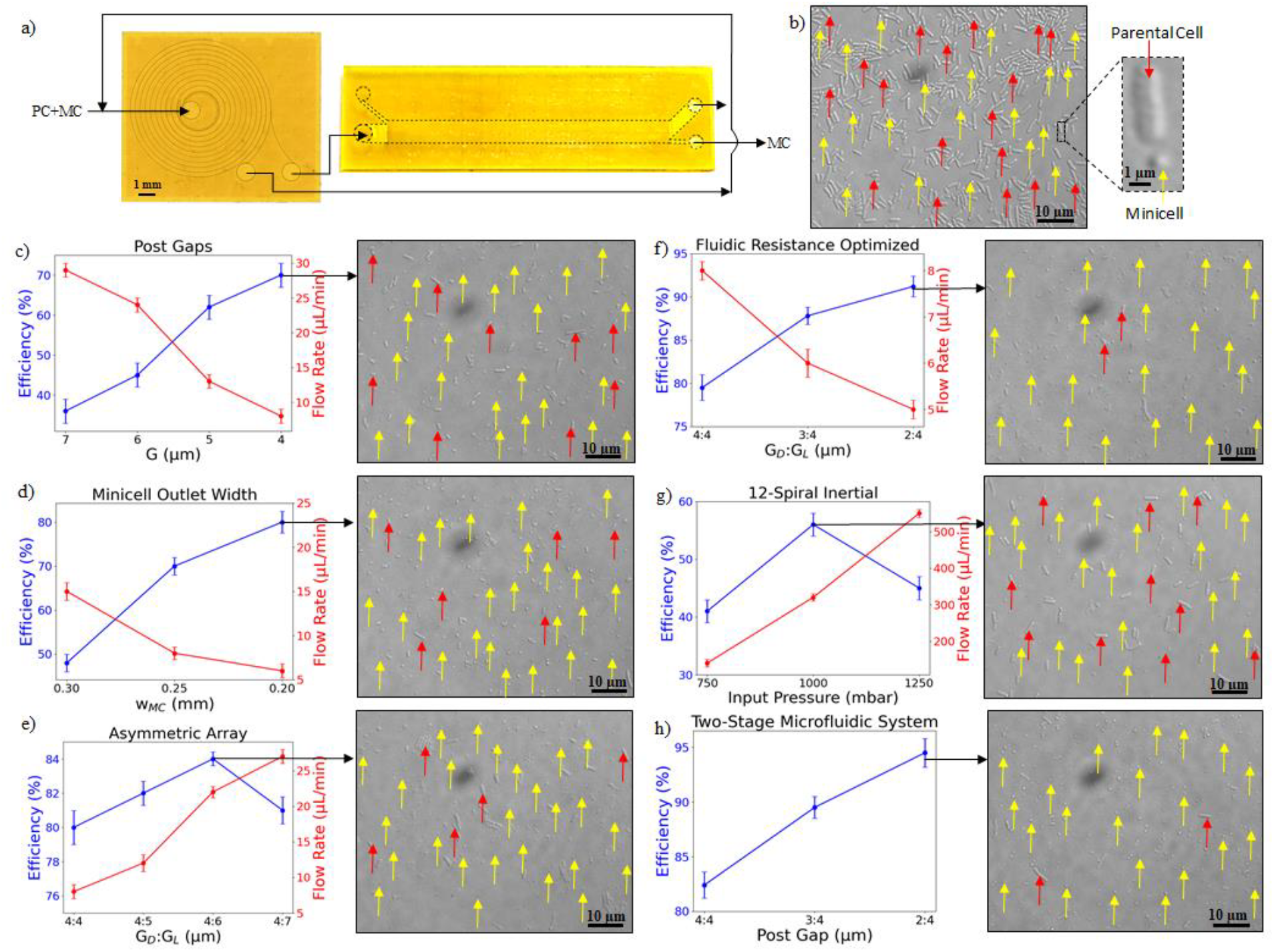
Varying lateral post gaps in asymmetric arrays impacts separation efficiency. a) Two-stage microfluidic system schematic. PC, parental cell. MC, minicell. b) Input sample with zoomed image of a parental cell (red arrow) and a minicell (yellow arrow); c) Post Gaps; d) Minicell outlet width; e) Asymmetric array; f) Fluidic resistance optimized; g) 12-Spiral inertial chip; h) Two-stage spiral inertial and DLD chip in combination. b-h) Scale bar, 10 µm. The graphs show minicell separation efficiency. Data represent mean ± standard error (n = 3)

## 3. Conclusion

In this work, we developed an optimized DLD microfluidic system fabricated via TPL for separating bacterial minicells from their parental cells. TPL offers submicron precision and helped overcome typical challenges of high-resolution and high-aspect ratio designs. We optimized DLD array parameters to reduce flow resistance and increase separation efficiencies. Asymmetric array designs reduced the risk of clogging due to long parental cells and maintained stability under high cell loads, while the optimized fluidic configuration reduced fluidic resistance and allowed for DLD designs with smaller post gaps. To further enhance the efficiency, we developed a two-stage microfluidic platform integrating a spiral inertial microfluidic as the first stage, followed by the optimized DLD chip as the second stage. This integrated system successfully achieved a minicell separation efficiency of 94 %. Future work can focus on enhancing the purification process by increasing throughput and improving separation efficiency to achieve thorough purification, eliminating any parental cell. This is crucial, as live bacterial cells could lead to undesirable side effects. For the viable future adoption of minicells as a drug carrier for cancer therapy, it is essential to further improve both throughput and purification efficiency.

## 4. Experimental Section

### Materials

2-Isopropanol (IPA), borosilicate glass slides 75 mm x 25 mm x 1 mm were purchased from (Carl Roth, Germany), 3-methacryloxypropyldimethylchlorosilane (MACS) were purchased from abcr (Germany), Borosilicate glass substrate holder 20 mm x 20 mm x 5 mm and UpPhoto were purchased from UpNano (Austria), Polytetrafluoroethylene (PTFE) tubes with inner diameter 0.85 mm and outer diameter 1.2 mm, and HPLC connectors were purchased from Bohlender (Germany).

### Substrate Functionalization

Glass slide were cleaned using isopropanol, distilled water, followed by drying with compressed N_2_. Plasma cleaning was performed at low pressure for 5 minutes to remove contaminants and enhance surface adhesion. The slides are completely immersing in a MACS solution inside a desiccator with nitrogen atmosphere for 60 minutes. Finally, the slide is cleaned again with isopropanol, distilled water, followed by drying with compressed N_2_.

### Two-Photon Lithography

TPL was performed using the NanoOne (UpNano GmbH, Austria) equipped with a 10 x immersion objective (NA 0.4, UPLXAPO10X, Olympus, Austria) in vat mode. The laser (80 MHz, 90 fs pulse, 780 nm wavelength) was focused through a cover glass into the vat containing the UpPhoto resin, and the focal point was maintained at a constant height above the glass slides, which in our case were MACS functionalised glass slides of 38 mm × 25 mm × 1 mm affixed with a double-sided tape on the provided glass substrate of 25 mm × 25 mm × 10 mm

### Fused Deposition Modelling

To facilitate high-pressure flushing of the microchannels, a Chip-to-World Interface (CWI) was fabricated using a Prusa i3 MK3S printer (Prusa Research, Czech Republic). The CWI was printed using polylactic acid (PLA), selected for its ease of processing and an infill of 25% for mechanical stability.

### High-Pressure Development

The CWI comprises two modular components that are secured together with screws and nuts. For an airtight connection, microfluidic tubing is clamped to the top section using a thermoelectric flanging tube (Bola, Germany). High-Performance Liquid Chromatography (HPLC) connectors, rated for high pressures, join the inlets and outlets to the chip. For channel development, isopropanol was pumped at 6.9 bar through the sealed microchannels using a microfluidic pump (Fluigent, France) connected to a main pressure supply. The pump’s outlet was connected to a conical tube (Eppendorf, Germany) with a P-Cap, while the chip outlets were directed into test tubes.

### Sample Preparation

To produce minicell, we transformed *E. coli* BL21(DE3) cells with pBLADE-minDΔ10.^[42]^ BLADE is a blue light-responsive transcription factor. Upon blue light induction, BLADE triggers the expression of a truncated MinD protein, which leads to minicell formation.^[42]^ The strain was additionally transformed with pUC19-pLac-sfGFP for constitutive superfolder green fluorescent protein (sfGFP) expression to facilitate fluorescence imaging. Overnight cultures grown from glycerol stocks in Luria–Bertani (LB) medium were diluted 1:100 into fresh LB and incubated at 37 °C with shaking till their OD_600_ reach 0.6. The incubation continued at 25 °C for 20 hours. The culture was then illuminated with blue light for 3 hours at 37°C using a custom-built LED device to trigger minicell formation. Current was fixed at 1.69 A using a regulated power supply; voltage and power varied accordingly (approx. 23 V and 36–37 W, respectively). Post-incubation, cells were washed twice and suspended in phosphate-buffered saline (PBS) solution.

### Flow Rate

The flow rates were measured using a flow rate sensor (Fluigent, France) connected to the microfluidic pump that is integrated with the OxyGEN software.

### Fluorescence Microscopy

Following microfluidic separation, output samples were collected and deposited on 2% agarose pads to immobilize cells for imaging. Fluorescence images were acquired using a Zeiss Axio Observer Z1/7 inverted wide-field fluorescence microscope (Carl Zeiss, Germany). sfGFP fluorescence was excited using a 38 HE filter set, which comprises an excitation filter (450–490 nm), a dichroic beam splitter, and an emission filter (500–550 nm). This configuration minimized background noise and allowed high-resolution imaging to assess minicell purity and yield. The parental cell and minicell numbers were counted manually from the microscope images.

## Acknowledgements

This project has received funding from the European Research Council (ERC) under the European Union’s Horizon 2020 research and innovation program (Grants Agreement No. 816006 to B.E.R. and No. 101002044 to B.D.V.), from the Deutsche Forschungsgemeinschaft (DFG), German Research Foundation) under Germany’s Excellence Strategy – EXC-2193/1 – 390951807 and EXS-2189-Project ID 390939984 (CIBSS). This work was conducted within the Research Cluster “Interactive and Programmable Materials (IPROM)” funded by the Carl Zeiss Foundation.

Received: ((will be filled in by the editorial staff)) Revised: ((will be filled in by the editorial staff)) Published online: ((will be filled in by the editorial staff))

## Supporting Information

**Figure S1.**
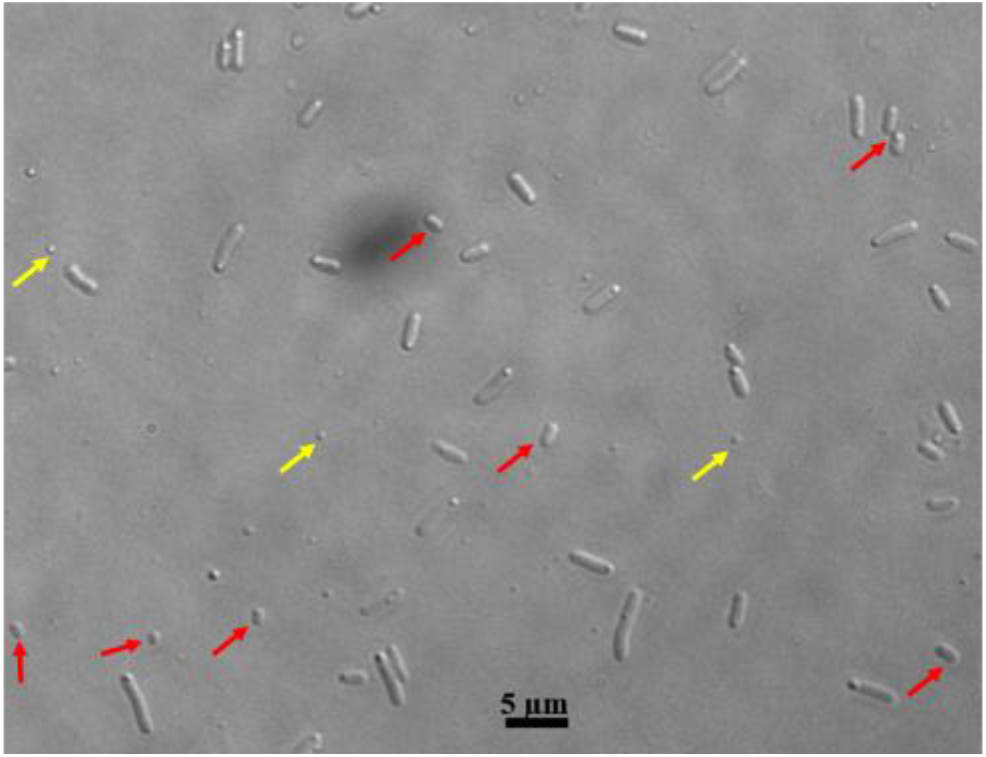
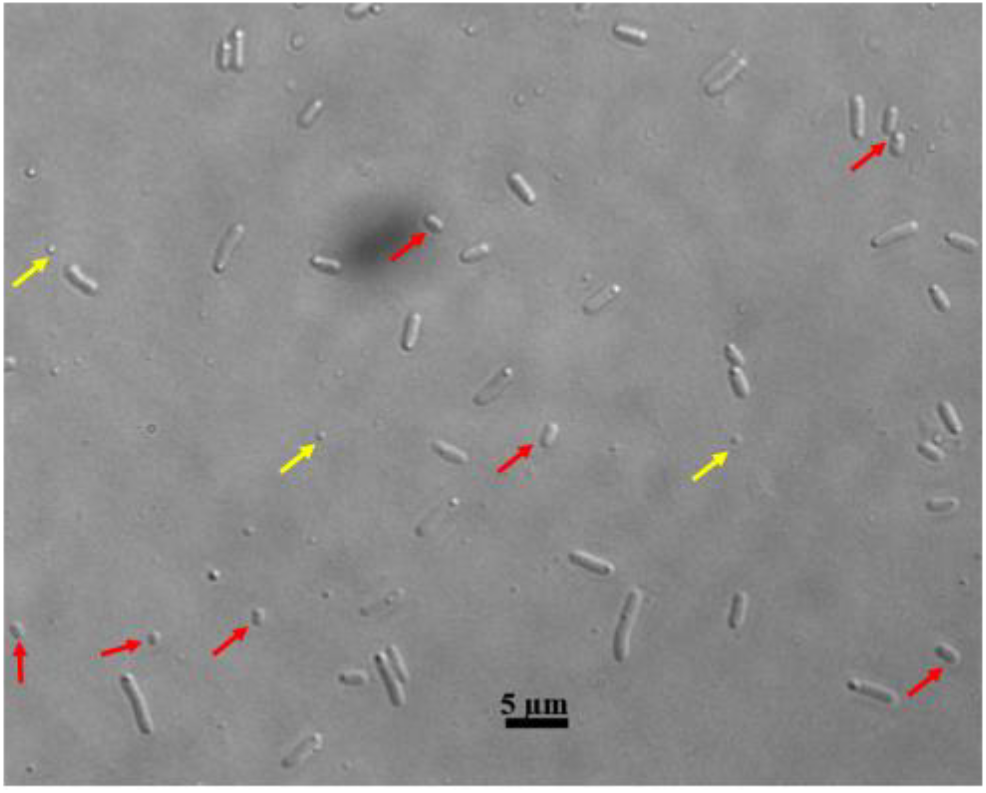
DIC microscopy image of the cell population after blue light illumination. Yellow arrows point to minicells, while red arrows highlight very short parental cells.

The surface of the glass slide is initially plasma cleaned (**Figure S1a)** and then functionalized with MACS (**Figure S1b)** before TPL printing. Following TPL printing, to develop microchannels and run the experiments, isopropanol is passed with an inlet pressure of 7 bar through the channels with a setup as seen in (**Figure S1c)**. A CWI ensures a seamless, airtight connection between the microfluidic chip and external tubing, preventing air gaps or leaks that could disrupt the process as seen in. This setup enables effective high-pressure flushing, eliminating undeveloped resin and ensuring unobstructed microchannels for optimal performance.

**Figure S2.**
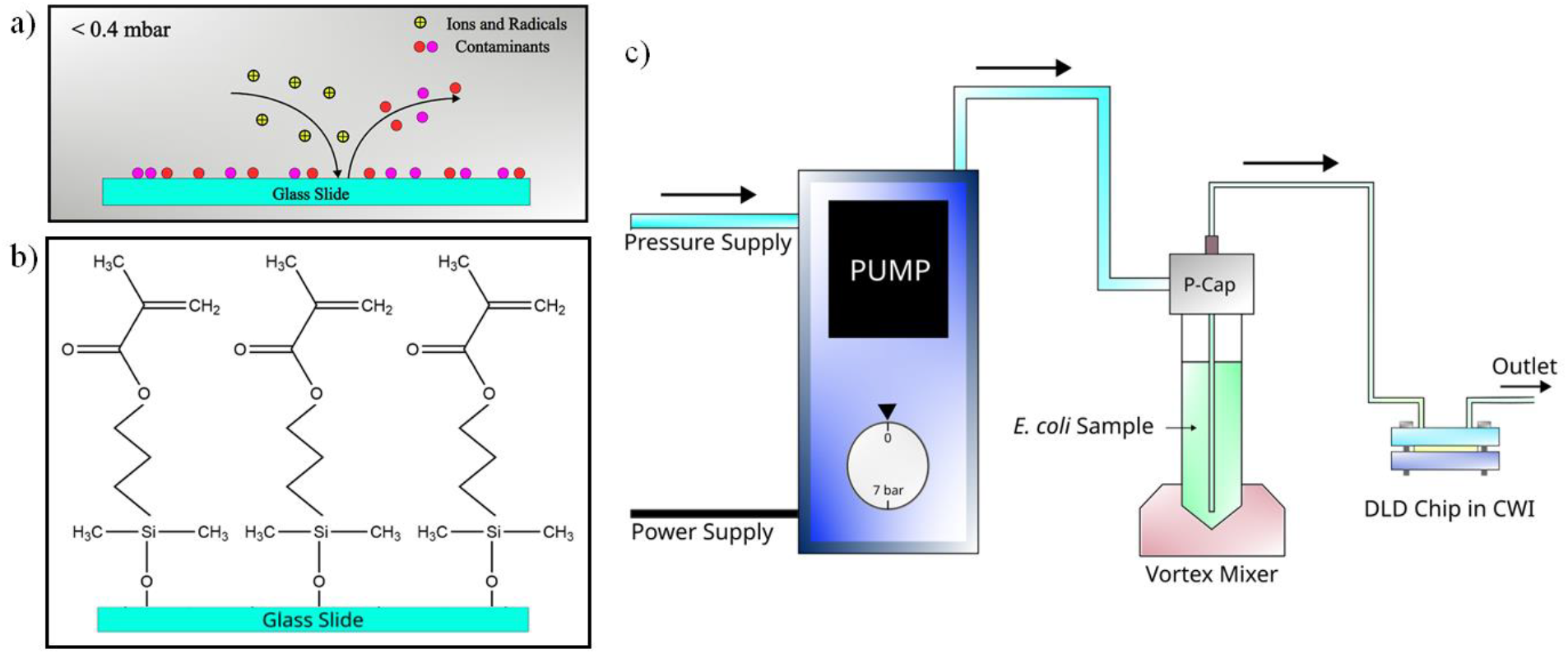
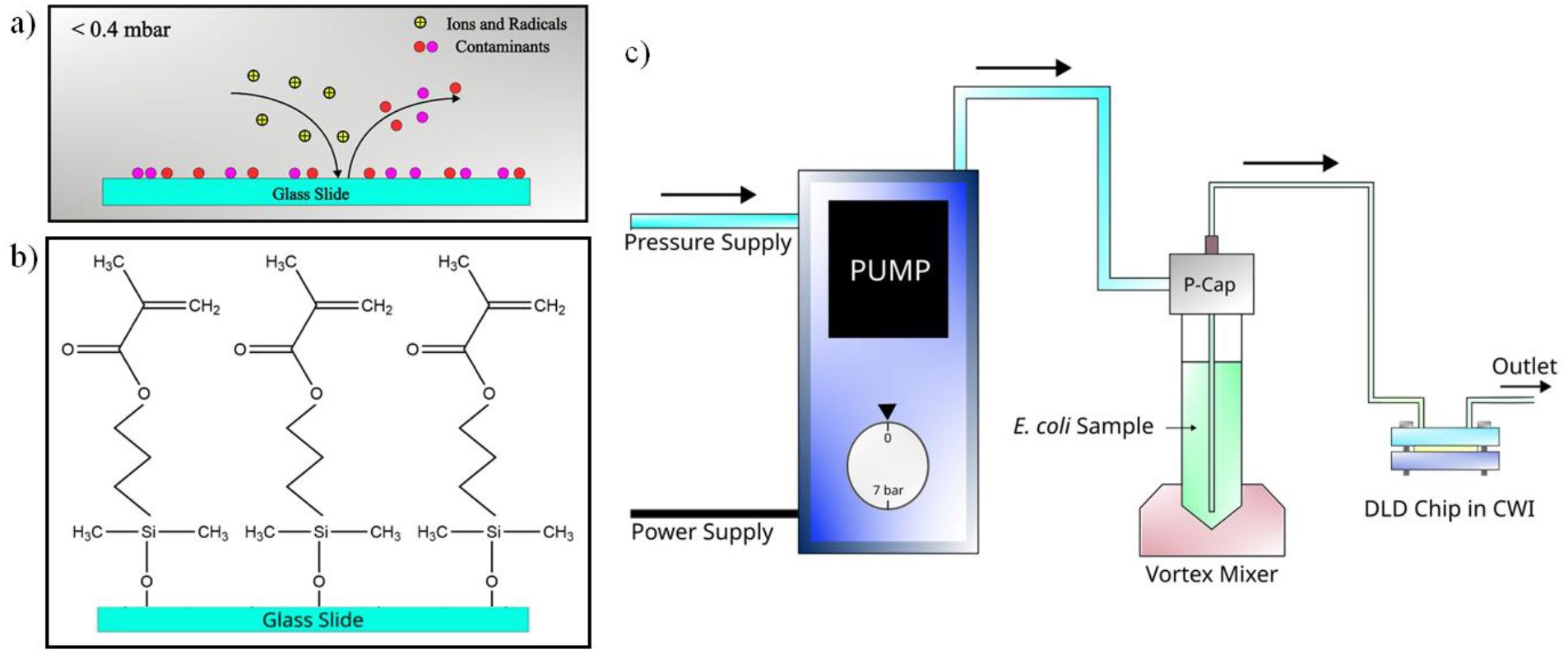
a) Plasma cleaning of the glass slide b) MACS functionalization of the glass slide c) Schematic of minicell purification experimental setup

**Table S1.**
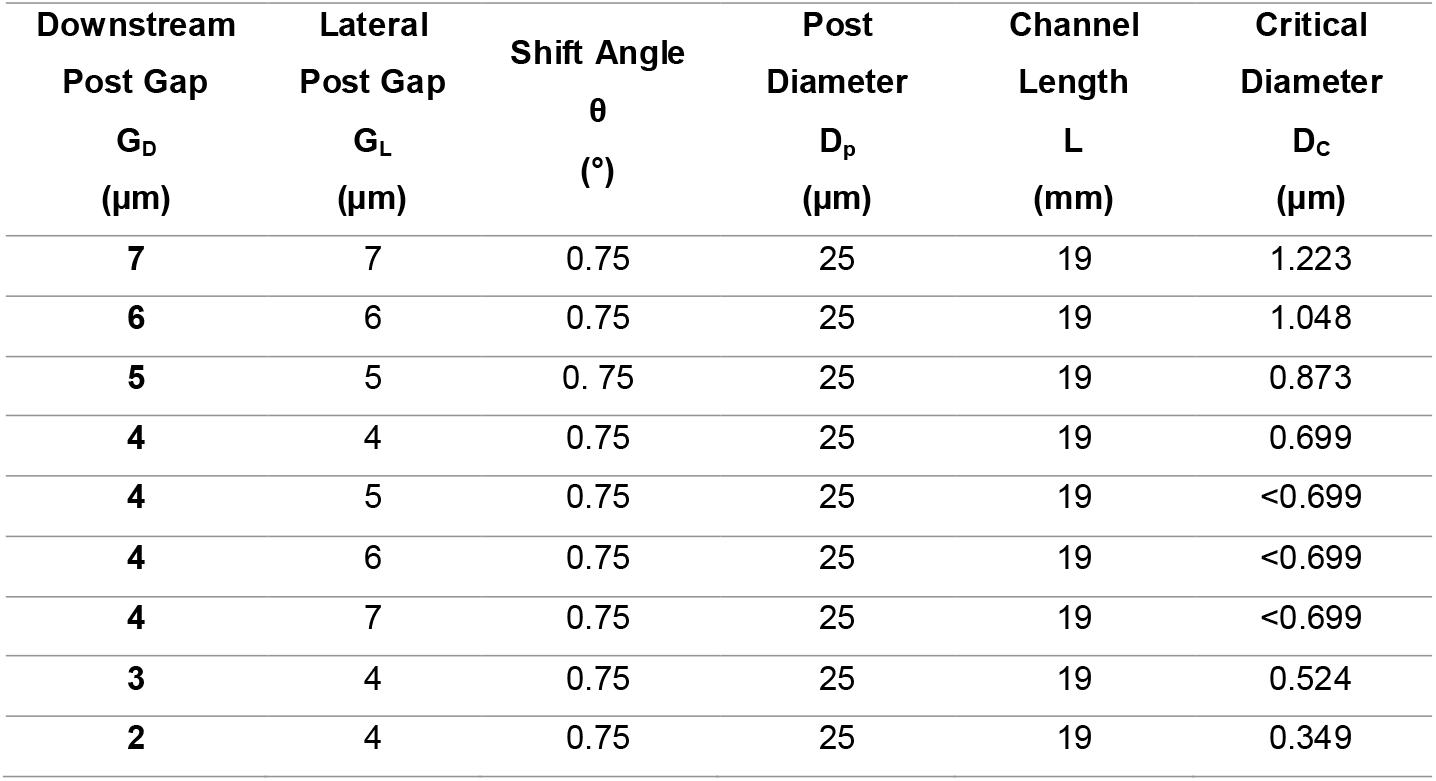
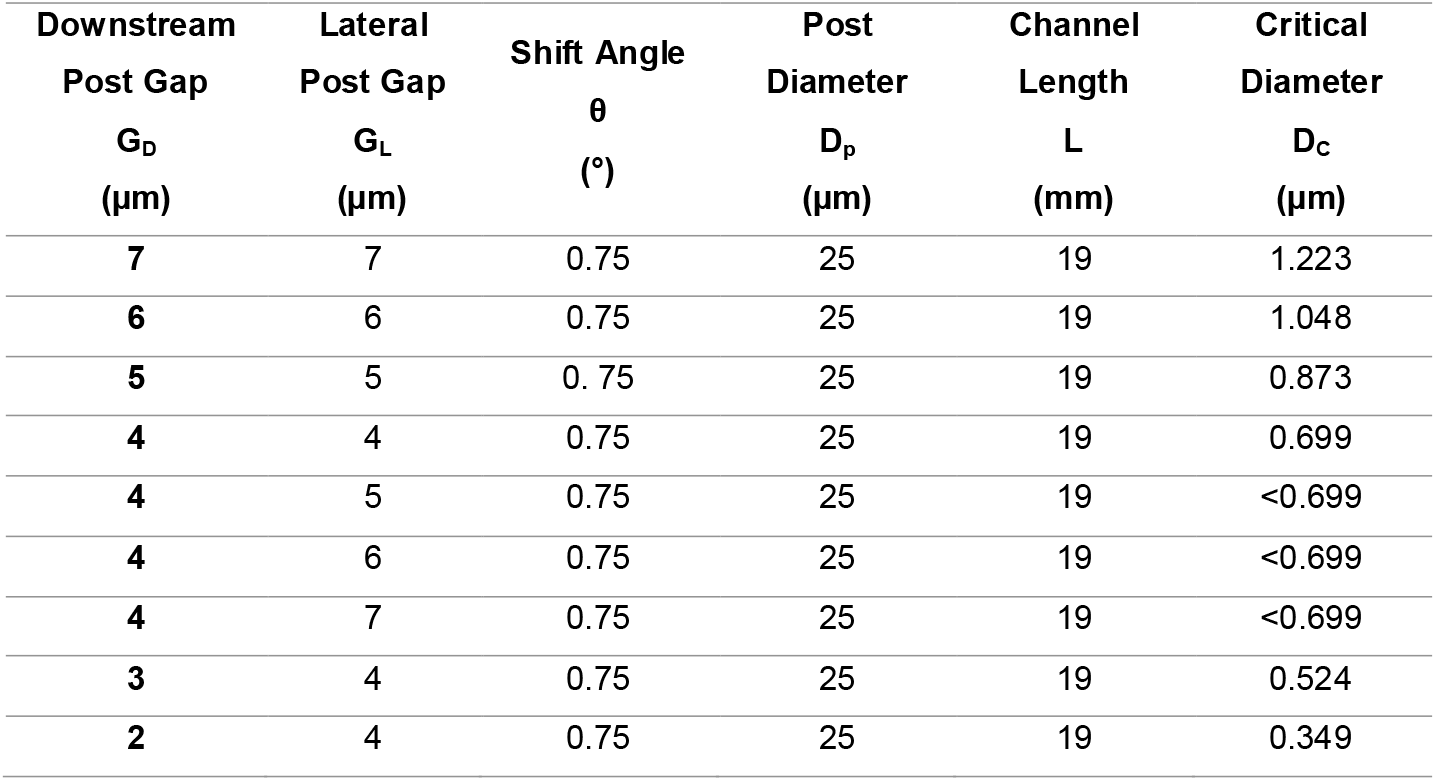
DLD parameters and resulting critical diameters.

| Downstream<br>Post Gap<br>$G_D$<br>($\mu\text{m}$) | Lateral<br>Post Gap<br>$G_L$<br>($\mu\text{m}$) | Shift Angle<br>$\theta$<br>($^\circ$) | Post<br>Diameter<br>$D_p$<br>($\mu\text{m}$) | Channel<br>Length<br>$L$<br>(mm) | Critical<br>Diameter<br>$D_c$<br>($\mu\text{m}$) |
| --- | --- | --- | --- | --- | --- |
| 7 | 7 | 0.75 | 25 | 19 | 1.223 |
| 6 | 6 | 0.75 | 25 | 19 | 1.048 |
| 5 | 5 | 0.75 | 25 | 19 | 0.873 |
| 4 | 4 | 0.75 | 25 | 19 | 0.699 |
| 4 | 5 | 0.75 | 25 | 19 | <0.699 |
| 4 | 6 | 0.75 | 25 | 19 | <0.699 |
| 4 | 7 | 0.75 | 25 | 19 | <0.699 |
| 3 | 4 | 0.75 | 25 | 19 | 0.524 |
| 2 | 4 | 0.75 | 25 | 19 | 0.349 |

